# Altered low frequency brain rhythms precede changes in gamma power during tauopathy

**DOI:** 10.1101/2021.08.03.454865

**Authors:** Fabio R. Rodrigues, Amalia Papanikolaou, Joanna Holeniewska, Keith G. Phillips, Aman B. Saleem, Samuel G. Solomon

## Abstract

Alzheimer’s disease and other dementias are associated with disruptions of electrophysiological brain activity, including low frequency and gamma rhythms. Many of these dementias are also associated with the malfunction of the membrane associated protein tau. Tauopathy disrupts neuronal function and the stability of synapses and is a key driver of neurodegeneration. Here we ask how brain rhythms are affected by tauopathy, at different stages of its progression. We performed local field potential recordings from visual cortex of rTg4510 and control animals at early stages of neurodegeneration (5 months) and at a more advanced stage where pathology is evident (8 months). We measured brain activity in the presence or absence of external visual stimulation, and while monitoring pupil diameter and locomotion to establish animal behavioural states. At 5 months, before substantial pathology, we found an increase in low frequency rhythms during resting state in tauopathic animals. This was because tauopathic animals entered intermittent periods of increased neural synchronisation, where activity across a wide band of low frequencies was strongly correlated. At 8 months, when the degeneration was more advanced, the increased synchronisation and low frequency power was accompanied by a reduction in power in the gamma range, with diverse effects across different components of the gamma rhythm. Our results indicate that slower rhythms are impaired earlier than gamma rhythms in tauopathy, suggesting that electrophysiological measurements can indicate both the presence and progression of tauopathic degeneration.

## Introduction

Several neurodegenerative disorders, including Alzheimer’s Disease (AD), are characterized as tauopathies because of the malfunction of the microtubule associated protein *tau*. In these tauopathies, tau is hyperphosphorylated, disrupting neuronal function (Roy et al., 2005; Crimins et al., 2011; Kopeikina et al., 2013) and impairing the stability of synapses (Hoover et al., 2010; Jackson et al., 2017). These impairments to neuronal structure and function should have correlates in the electrical activity of the brain, so the electroencephalogram (EEG) and related measurements could offer potentially powerful, low-cost tools for detecting and tracking the functional impact of tauopathy.

Neuronal activity often coheres into brief rhythms (Buzsaki and Draguhn, 2004). EEG measurements from patients with AD indicate an increased power at low frequencies (delta, ca. 2-4Hz) (Coben et al., 1983; Huang et al., 2000), and reduced power at higher frequencies including alpha, beta and gamma-bands (for a review see Babiloni et al. (2020)). The contribution of tauopathy to these changes is unclear. In preclinical mouse models of tauopathy, low frequency EEG oscillations are increased (Das et al., 2018), fMRI resting state networks are disrupted (Green et al., 2019), and coupling between hippocampal and prefrontal cortical local field potentials (LFPs) is disrupted (Ahnaou et al., 2017). In the well-characterized rTg4510 model of tauopathy, hippocampal LFP (Ciupek et al., 2015) and cortical EEG are disrupted (Holton et al., 2020), and strong low frequency oscillations emerge in LFP recordings from frontal cortex (Menkes-Caspi et al., 2015). (Menkes-Caspi et al., 2015). Limited work has characterized gamma-band oscillations (30-120Hz), and found reduced gamma in the hippocampus (Booth et al., 2016a) and entorhinal cortex (Booth et al., 2016b) of rTg4510 mice. However, it is not clear if changes in oscillations at low and high frequencies occur together, or if one precedes the other.

Frontal cortex and hippocampal areas are associated with complex cognitive functions (e.g. motivation and memory), that make it difficult to make controlled measurements in awake animals. Sensory brain areas, such as visual cortex, offer the opportunity to characterise the influence of tauopathy on brain activity during presence and absence of external stimulation (visual stimulus), while monitoring behavioural states such as arousal and locomotion (Niell and Stryker, 2010; Reimer et al., 2014; Vinck et al., 2015). EEG and LFP signals from the mouse primary visual cortex (V1) are dominated by low frequency oscillations (2-6Hz) (Senzai et al., 2019) with additional oscillations in the gamma range (Buzsaki and Draguhn, 2004), including components driven by thalamic inputs (Saleem et al., 2017; Storchi et al., 2017; McAfee et al., 2018).

In this study, we evaluated the influence of tauopathy on electrical activity in visual cortex by making recordings from V1 in the rTg4510 mouse model. Recordings were made at 5 months, early in neurodegeneration, or at 8 months, when neurodegeneration is more advanced (Ramsden et al., 2005; Santacruz et al., 2005). We found a pronounced increase in low frequency oscillations early in tauopathy, which was linked to the occurrence of abnormal periods of high cortical synchronization. Gamma power and stimulus-induced gamma oscillations were reduced only at more advanced stages of neurodegeneration. We conclude that increased low frequency rhythms may be an early marker of tauopathy and associated disorders.

## Results

We measured the local field potential (LFP) from electrodes chronically implanted into layer 4 of V1 of awake rTg4510 animals and their WT littermates. At 2 months of age, half the rTg4510 animals were transferred to a doxycycline-containing diet to suppress the expression of the transgene (henceforth Tau- animals), while the remaining animals were maintained on a normal diet (Tau+) (Fig 1A). As expected (Santacruz et al., 2005), phosphorylated-tau burden (as indicated by staining for AT8 antibody) was higher in 5m and 8m Tau+ animals than in Tau- or WT animals (Fig 1B). The brain weight of Tau+ and Tau- animals was similar at 5m, and both were smaller than those of WT littermates, but at 8m brain weight of Tau+ animals was smaller than that of Tau- animals, indicating degeneration between 5m and 8m (Fig 1C).

**Figure 1 –.**
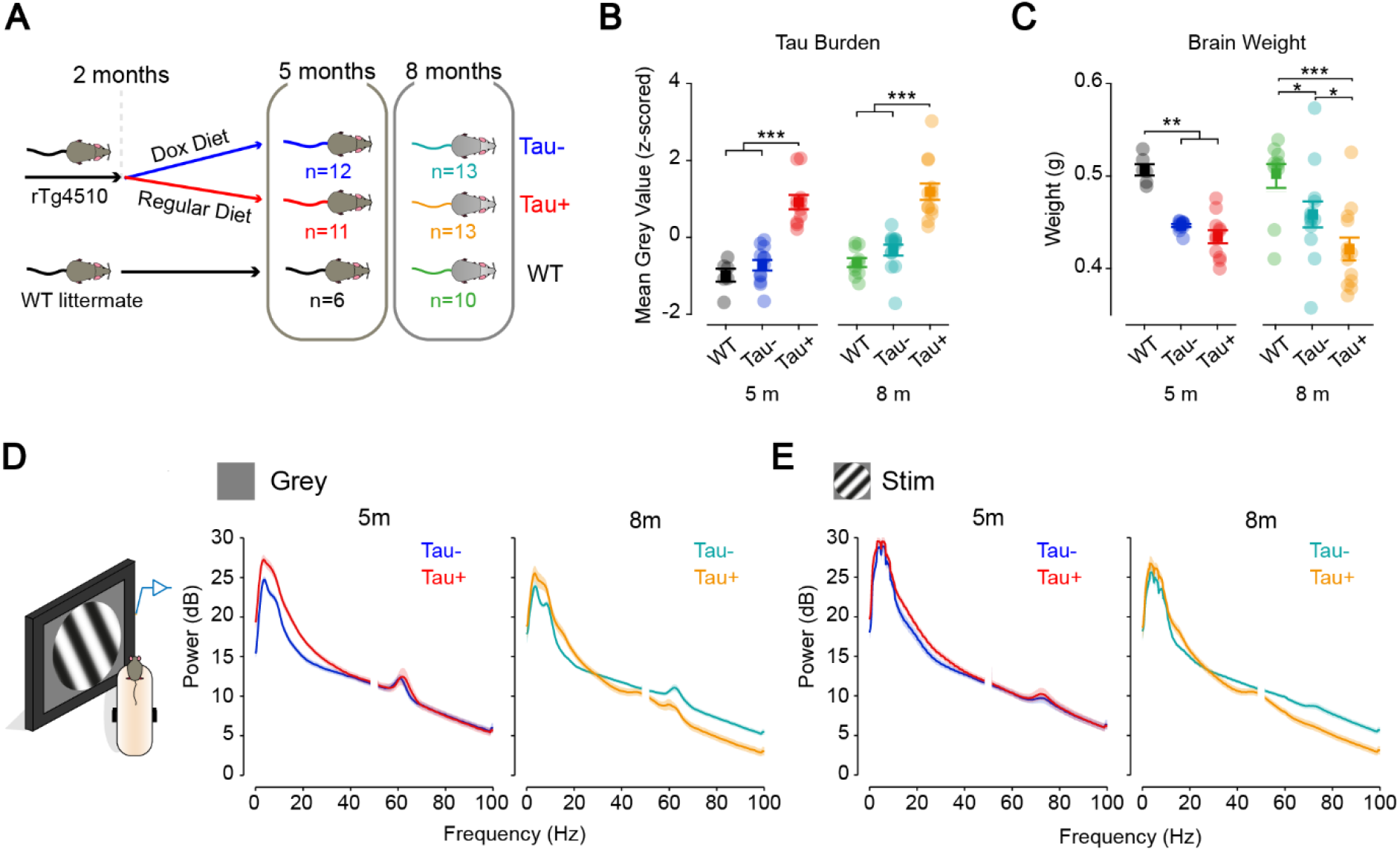
Visual cortical oscillations are disrupted in the rTg4510 mouse model. (A) At 2 months, 24 Tg4510 mice were maintained on a regular diet and continued to express human mutant tau (Tau+) while another 25 animals were switched to a doxycycline-containing diet to arrest mutant tau expression (Tau-). WT littermates were used as controls for the experiment. Mice were tested at one of two different ages: 5 months (5m), before the onset of neurodegeneration; and 8 months (8m), at more advanced stages of degeneration. (B) The levels of phosphorylated human tau were quantified for each mouse group following immunostaining. Quantification of tau burden (by measuring average pixel value) showed significantly higher levels of tau present in Tau+ mice (*F*_(2,2)_=947.4) at both tested ages (5m, 0.914±0.187a.u.; 8m, 1.185±0.213a.u.) compared to Tau- (5m: −0.726±0.140a.u., *P*=8.91×10^-9^; 8m: −0.332±0.142a.u., *P*=1.35×10^-8^) and WT mice (5m, −0.9184±0.166a.u.; 8m, −0.654±0.117a.u.). (C) At earlier stages of tauopathy (5m), there is no difference in the brain size of Tau+ (0.434±0.007g) and Tau- (0.446±0.002g, *F*_(2,2)_=25.746, *P*=1.00) animals. At 8m, following degeneration, the mean brain size of Tau+ mice was smaller (8m, 0.421±0.012g) compared to Tau- animals of the same age (0.459±0.014g, *P*=0.030). WT animals at 5m (0.507±0.006g) and 8m (0.500±0.013g) have larger brains compared to both Tau- (5m, *P*=0.004; 8m, *P*=0.023) and Tau+ mice (5m, *P*=0.001; 8m, *P*=7×10^-6^. (D and E) LFP recordings from visual cortex were obtained from head-fixed mice exposed to blocks of visual stimulus (full field sinusoidal gratings that reversed contrast at a 1Hz frequency) interspaced by 30s grey screen intervals. We generated power spectra from those recordings for both grey screen (D) and stimulus presentation (E) periods. Low frequency (2-10Hz) oscillations in the visual cortex of Tau+ mice were increased in power in the absence of any stimulation, at 5m and 8m. A reduction in overall gamma power was observed at 8m, both in the presence and absence of any visual stimulation. Panels A to C are adapted from Papanikolaou et al. (2021).

### Visual cortical oscillations are disrupted in rTg4510 model of tauopathy

We measured LFP from head-fixed animals over seven consecutive days. On each day the animal was initially exposed to a uniform grey screen, after which we interleaved presentations of a large, flickering grating with additional presentations of the grey screen. We subjected the recording on each day to multitaper analysis (see Methods), which provided estimates of the power spectrum of the LFP at a resolution of 1s. We then collapsed data across epochs of grey screen or stimulus presentation, and averaged observations across days, to produce two estimates of the power spectrum in each animal.

The power spectrum of visual cortical LFP was altered in both early and more advanced stages of tauopathy. We first analysed LFP signals during the presentation of a grey screen. We found a pronounced increase in low frequency activity (frequencies up to about 10 Hz) in 5m and 8m old Tau+, when compared to Tau- (Fig 1D) and WT animals (not shown). At 8m, increased low frequency activity in Tau+ was also accompanied by a decrease in the high gamma-band activity (55-100Hz). During periods of visual stimulus presentation (Fig 1E), power at low frequency was similar between Tau+, Tau- and WT (not shown) animals, at both 5m and 8m. Because the stimulus was flickering at a rate of 1 Hz, it elicited strong visually-evoked potentials in all animals (see Papanikolaou et al. (2021)), and it is therefore difficult to distinguish the contribution of stimulus-induced power changes, from those associated with ongoing cortical activity. Visual stimulation at 8m did not, however, restore gamma band activity in Tau+ animals, suggesting that the power reduction at high frequencies cannot be rescued by visual stimulation.

### Impact of tauopathy on visual cortical activity is more prominent during resting state

Measurements of cortical electrical activity in AD patients show larger disruptions during periods of quiet rest (de Haan et al., 2008; Babiloni et al., 2009). We therefore asked if the changes in spectral power that we observed in the Tau+ animals were also more prominent during rest. During our measurements, animals were able to move freely over a styrofoam wheel. We recorded the animals’ movement and pupil diameter to monitor the behavioural state of the animal. From these measurements we defined moving, intermediate, and resting states (Fig 2A): ‘resting’ state was when the animal’s speed was less than 5cm/s, and the pupil size was smaller than the mean size over that recording session; ‘intermediate’ state was when the animal’s speed was less than 5cm/s and the pupil size was larger than the mean size over that recording session; ‘moving’ state was when the animal’s speed was greater than 5cm/s. We generated power spectra across moving, intermediate, and rest periods each day, in presence and absence of visual stimulus.

**Figure 2 –.**
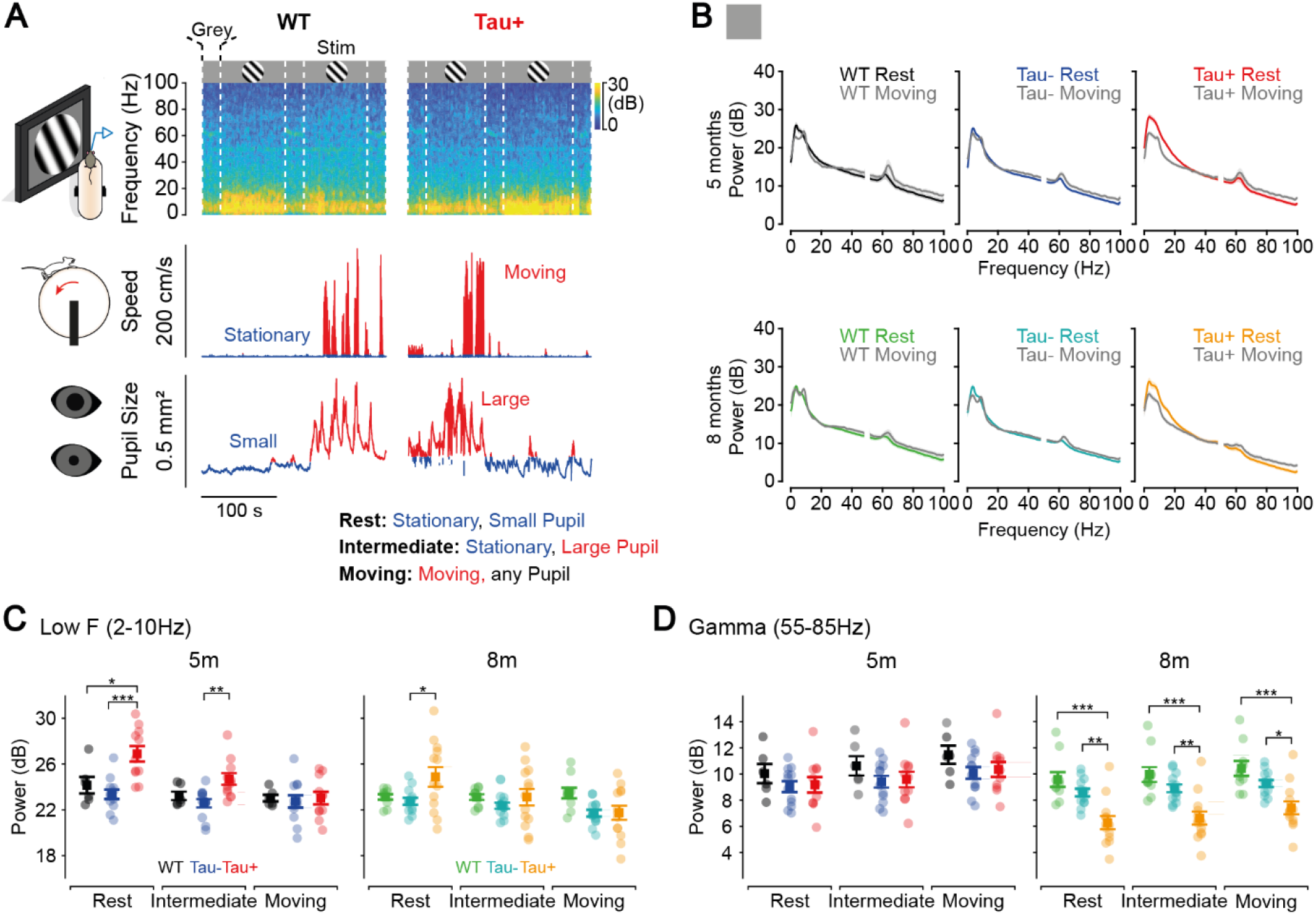
Differences in spectral power at low and gamma frequencies between Tau+ mice and controls are more pronounced during Rest periods. (A) Head-fixed animals, free to run over a Styrofoam wheel, were exposed to 10 blocks of 200 phase reversals of an oriented grating (which reversed at a frequency of 1 Hz), separated by grey screen intervals of 30s across 7 days. Speed of locomotion and pupil diameter were used to identify periods when the animal was stationary (speed<5cm/s) and when the pupil was small (pupil size<mean pupil size for recording session). Periods when the animal’s pupil was small, the animal was stationary, and no stimulus was presented were considered ‘Rest’ periods. ‘Intermediate’ periods where when the animal was stationary, and the pupil was large (pupil size>mean pupil size for recording session). ‘Moving’ periods where when the animal’s speed was above 5cm/s, regardless of pupil size. (B) During grey screen intervals, power at low frequencies was larger during rest periods compared to moving periods, particularly in Tau+ animals at both 5m and 8m. During moving periods, power in the gamma range is increased compared to rest periods, and this effect is similar across all groups and both age cohorts. (C) Average power during grey screen intervals at low frequencies (2-10Hz) for all animal groups and age cohorts during Rest, Intermediate and Moving periods was quantified. Low frequency power during grey screen presentations was significantly larger in Tau+ during Rest periods (5m, 26.873±0.679dB; 8m, 24.864±0.854dB) compared to controls (WT 5m, 24.132±0.724dB; WT 8m, 23.110±0.23dB; Tau- 5m, 23.337±0.415dB; Tau- 8m, 22.734±0.307dB; *F*_(2.489,59)_=15.039; WT x Tau+ 5m, *P*=0.022; Tau- x Tau+ 5m, *P*=1.58×10^-4^; WT x Tau+ 8m, *P*=0.108; Tau- x Tau+ 8m, *P*=0.021). Intermediate states caused a reduction in mean low frequency power across all animals (WT 5m, 23.204±0.371dB; WT 8m, 23.103±0.246dB; Tau- 5m, 22.556±0.366dB; Tau- 8m, 22.341±0.271dB; Tau+ 5m, 24.682±0.502dB; Tau+ 8m, 23.093±0.726dB). However, differences were still detected between Tau+ and Tau- mice, but only at 5m of age (Tau- x Tau+ 5m, *P*=0.006). During moving periods, low frequency power in Tau+ animals was similar to controls (WT 5m, 23.008±0.291dB; WT 8m, 23.441±0.462dB; Tau- 5m, 22.716±0.541dB; Tau- 8m, 21.689±0.299dB; Tau+ 5m, 23.014±0.543dB; Tau+ 8m, 21.729±0.619dB). (D) Mean high gamma (55-85Hz) power was similar between Tau+ and control groups at 5m during rest (WT, 10.015±0.729dB; Tau-, 9.015±0.418dB; Tau+, 9.155±0.592dB), intermediate (WT, 10.546±0.742dB; Tau-, 9.329±0.450dB; Tau+, 9.501±0.600dB) and moving periods (WT, 11.442±0.699dB; Tau-, 10.056±0.448dB; Tau+, 10.315±0.567dB). At advanced stages of degeneration, mean gamma power in Tau+ during was smaller than that of controls during rest (WT, 9.569±0.557dB; Tau-, 8.568±0.294dB; Tau+, 6.266±0.502dB; *F*_(2.845,59)_=3.979; WT x Tau+ 8m, *P*=3.4×10^-5^; Tau- x Tau+ 8m, *P*=0.002), intermediate (WT, 9.939±0.571dB; Tau-, 8.879±0.301dB; Tau+, 6.596±0.499dB; WT x Tau+ 8m, *P*=4×10^-5^; Tau- x Tau+ 8m, *P*=0.003), and moving periods (WT, 10.399±0.569dB; Tau-, 9.240±0.265dB; Tau+, 7.383±0.488; WT x Tau+ 8m, *P*=1.20×10^-4^; Tau- x Tau+ 8m, *P*=0.014).

The increase in low frequency oscillations in Tau+ animals was more pronounced during rest (Fig 2B, C). The spectral power at low frequencies (averaged over 2-10Hz) was ~13% larger in Tau+ at 5m than in control animals at rest, and ~8% larger at 8m. In contrast, there were no differences in low frequency power between Tau+ and control animals during moving periods (Fig 2C). Intermediate states were accompanied by smaller reductions in low frequency power for all animal groups at 5m (8% reduction in Tau+, 4% in Tau- and 3% in WT), resulting in the mean power in Tau+ remaining larger than that in Tau- animals, but not WT. At 8m there were no differences in the mean low frequency power during intermediate states. We showed that presentation of flickering visual stimuli abolished differences in low frequency power between Tau+ and Tau- animals (Fig 1) – this was also the case when we considered moving, intermediate, and resting periods separately (not shown).

We found a significant increase in gamma power during moving and intermediate state periods relative to rest, in all animals and ages (Fig 2D). Despite this state-dependent increase in power, gamma in 8m Tau+ animals was consistently lower than that of control animals, regardless of behavioural state.

### Late tauopathy disrupts high gamma-band activity

Gamma-band activity in the visual cortex can be decomposed into multiple sub-bands (Engel et al., 2001; Chen et al., 2017; Veit et al., 2017). Activity at some frequencies (e.g., ‘narrow-band gamma’; ca. 55-65 Hz; (Saleem et al., 2017)) may arise by entrainment of visual cortex to oscillations that are already present in the thalamocortical input (Saleem et al., 2017; Storchi et al., 2017; McAfee et al., 2018). Activity in other sub-bands may be more reliant on cortico-cortical signals, or reflect the activity of local interneurons (e.g., parvalbumin interneuron driven oscillations at ca. 70-80Hz) (Veit et al., 2017). We showed that overall gamma-band activity is reduced in 8m Tau+ animals, during presentation of both grey screen and visual stimulus (Fig 1). However, the power spectrum in the high gamma-band range also showed clear ‘bumps’, and the peak frequencies of these bumps were different in the grey screen and stimulus conditions (Fig 3A). As expected from previous work (Saleem et al., 2017), gamma power peaked in a narrow range around 55-65Hz during the grey screen condition. During the presentation of the visual stimulus, this peak disappeared and another emerged at higher frequencies, around 70-80Hz (Veit et al., 2017; McAfee et al., 2018). To characterise these gamma peaks we first calculated the aperiodic component of the power spectra, using standard methods (Donoghue et al., 2020) (Fig 3A). The aperiodic component decreases monotonically as a function of frequency, and we subtracted (in dB) this aperiodic component from the raw spectrum to produce a normalized spectrum (Fig 3B).

**Figure 3 –.**
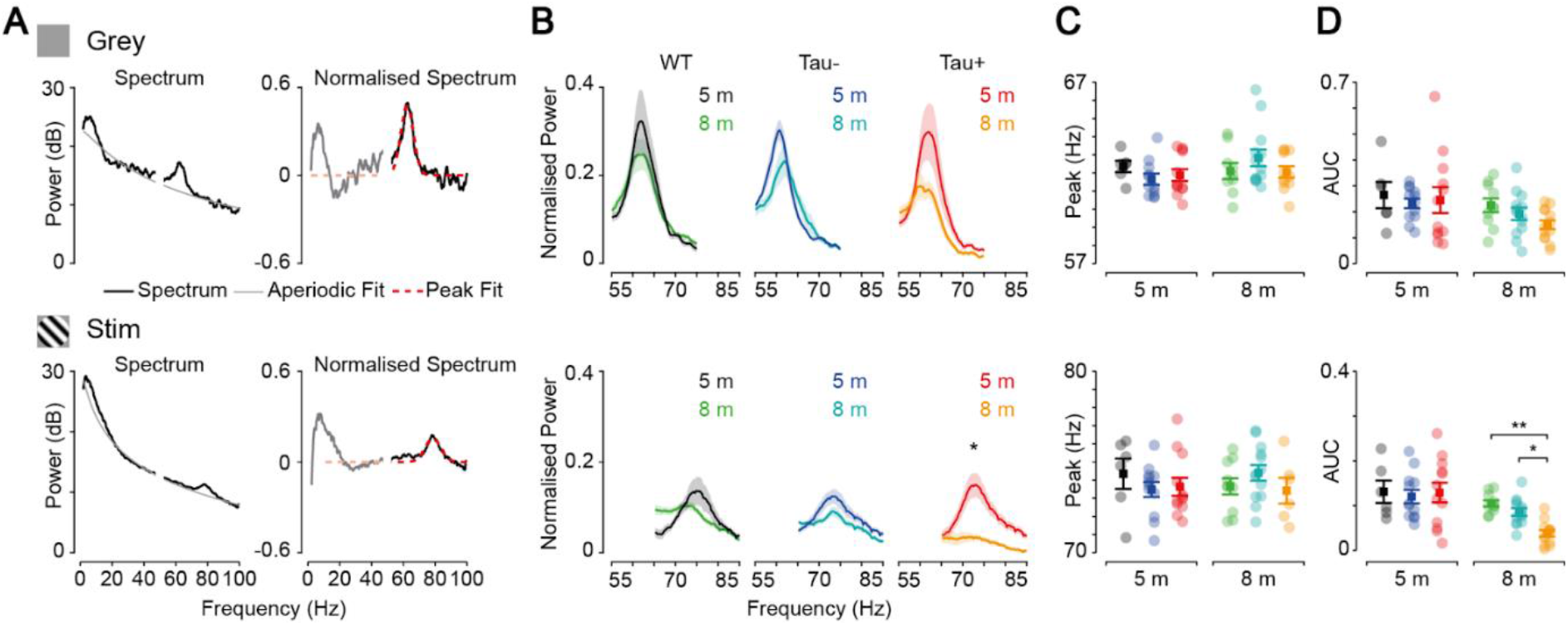
Stimulus-induced gamma oscillations are reduced in late neurodegeneration. (A) The power spectra of all animals were normalised by dividing the spectra by their corresponding the aperiodic component (1/f trend line). These normalised spectra were used to identify gamma oscillations during grey screen and stimulus presentation. (B) Average normalised spectrum during rest periods for all mouse groups and age cohorts at frequency ranges where grey- and stimulus-evoked oscillations are detected (55-85Hz) are shown. Gamma oscillations amplitudes tended to decrease with age both during grey (WT 5m, 0.265±0.051dB; WT 8m, 0.225±0.027dB; Tau- 5m, 0.233±0.019dB; Tau- 8m, 0.192±0.024dB; Tau+ 5m, 0.260±0.054dB; Tau+ 8m, 0.150±0.017dB), and stimulus presentation (WT 5m, 0.131±0.025db; WT 8m, 0.105±0.007dB; Tau- 5m, 0.120±0.015dB; Tau- 8m, 0.086±0.009dB; Tau+ 5m, 0.137±0.023dB; Tau+ 8m, 0.034±0.007dB). Neurodegeneration only significantly impacted the amplitude stimulus-induced gamma oscillation (*F*_(2,59)_=3.976; *P*=3×10^-6^). (C) Centre frequency of detected peaks in the respective gamma ranges during and grey (WT 5m, 62.355±0.313Hz; WT 8m, 62.106±0.441Hz; Tau- 5m, 61.652±0.307Hz; Tau- 8m, 62.831±0.461Hz; Tau+ 5m, 61.741±0.325Hz; Tau+ 8m, 62.055±0.313Hz) and stimulus presentation (WT 5m, 74.347±0.834Hz; WT 8m, 73.659±0.447Hz; Tau- 5m, 73.483±0.413Hz; Tau- 8m, 74.400±0.430Hz; Tau+ 5m, 73.707±0.535Hz; Tau+ 8m, 73.419±0.712Hz) did not significantly change with aging or tau load (*F*_(2,55)_=1.638). (D) The area under the curve (AUC) of a 6Hz range centred in the mean frequency values calculated in C, showed a significant reduction in AUC during stimulus-evoked gamma oscillations in Tau+ mice at 8m (WT x Tau+, *P*=0.004; Tau- x Tau+, *P*=0.033), but not at 5m or during grey screen periods (*F*_(2,59)_=0.840).

Gamma-bumps were present during presentation of grey screen and visual stimuli in all groups of animals, but the latter were more susceptible to tauopathy. We extracted the peak frequency of the best-fitting Gaussian for each condition in each animal, and found that the peak frequencies during grey screen, or during stimulus presentation, did not vary with animal group (Fig 3C). To compare the amplitude of activity around these peaks we calculated the area-under-the-curve (AUC, or amplitude) of the normalized power spectra in a 6Hz range centred on the mean peak frequency for each animal group. The amplitude of grey gamma-bumps was similar across age and groups. By contrast, the amplitude of stimulus gamma-oscillations reduced from 5m to 8m in all groups, particularly in Tau+ animals, such that the AUC was smaller in 8m Tau+ than in 8m Tau- or WT animals. We conclude that stimulus induced gamma oscillations are more susceptible to degeneration than the narrowband gamma oscillations that occur in the absence of a visual stimulus.

### Visual cortical ensembles are synchronised during tauopathy

We showed that low frequency power is increased in Tau+ animals during rest. Inspection of individual spectrograms further showed that the resting state in Tau+ animals (Fig 4A) was organised into epochs during which there was a synchronised increase in power across a wide band of low frequencies. The LFP traces in these epochs were characterized by strong and coherent low frequency oscillations (Fig 4A). Spectrograms obtained from Tau- and WT animals showed a more variegated structure of power at low frequencies (Fig 4A) and lacked the clear transitions into and out of these strongly synchronised states. To characterise the fluctuations in low frequency power we calculated a synchronisation (‘sync’) index as the ratio of low frequency (210Hz) to high gamma (55-85Hz) power in each time bin. A higher sync-index indicates relatively stronger power in the low frequency range.

**Figure 4 –.**
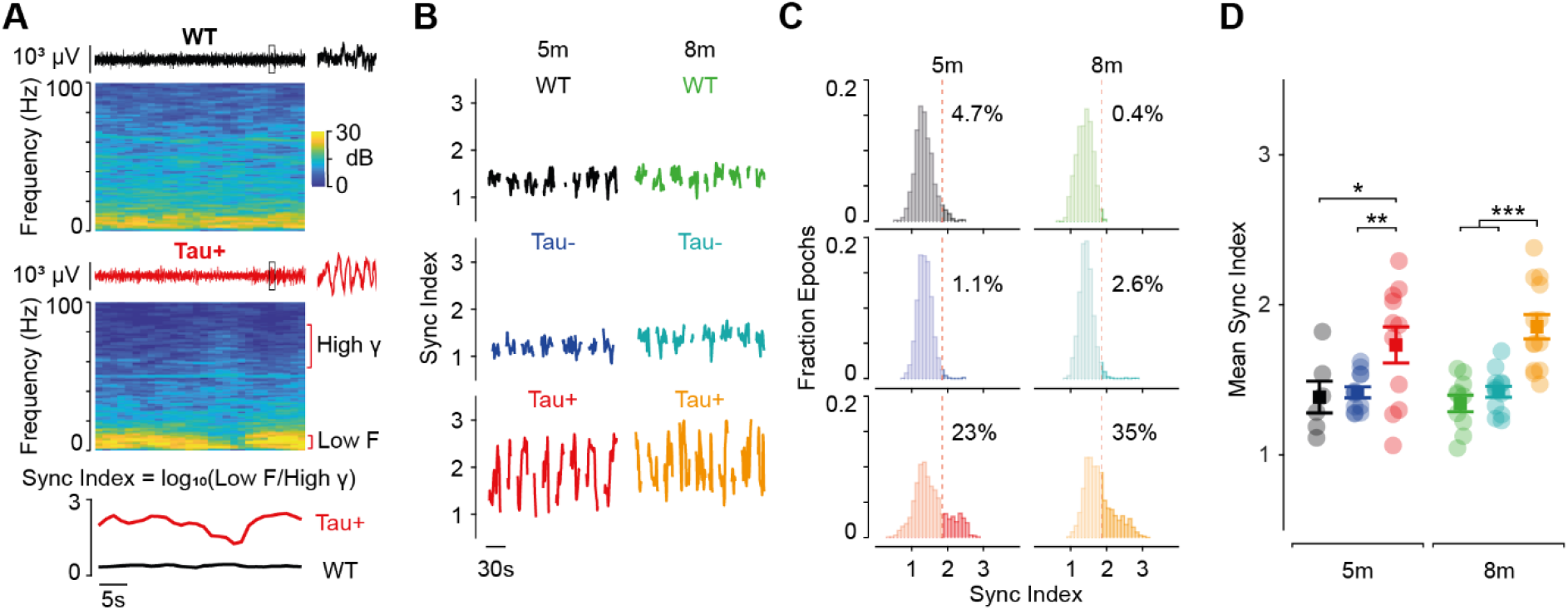
Increased ‘synchronization’ of visual cortical activity during tauopathy. (A) Example spectrograms for a grey screen interval and respective LFP trace for a WT and a Tau+ animal. The inset on the LFP trace corresponds to a 1s period highlighted in the trace. We observed increased modulation of low frequency oscillations in Tau+ animals relative to controls, and these were associated with larger low frequency oscillations in the LFP. We defined a ‘synchronization (sync) index’ of the activity in visual cortex at rest as the logarithm of the ratio between low frequencies (2-10Hz) and high gamma (55-85Hz) power, where larger sync index values reflect larger low frequency power relative to high gamma. (B) In Tau+ animals, visual cortex entered periods of increased synchronization (as shown by larger oscillations of this index), both in early and advanced degeneration. (C) The distribution of the sync index over all rest periods for all recording sessions also reflects the presence of periods of increased synchronization in Tau+ animals. We defined a sync threshold as the 95th percentile of the pooled WT distributions from both ages (red dashed line, 1.855). There were more instances of higher synchronization in Tau+ animals (>20% at both ages), than controls. (D) The mean sync index was larger for Tau+ animals during rest periods at both ages (WT 5m, 1.384±0.106; WT 8m, 1.342±0.056; Tau- 5m, 1.416±0.036; Tau- 8m, 1.419±0.037; Tau+ 5m, 1.732±0.121; Tau+ 8m, 1.854±0.081; *F*_(4.411,130.139)_=6.645; WT 5m x Tau+ 5m, *P*=0.023; Tau- 5m x Tau+ 5m, *P*=0.009; WT 8m x Tau+ 8m, *P*=3.3×10^-5^; Tau- 8m x Tau+ 8m, *P*=1.49×10^-4^).

We found high and variable sync-index in Tau+ animals, with epochs of strong synchronisation interspersed by epochs of weak synchronisation (Fig 4B); these strongly synchronised states were more prominent in some Tau+ animals than in others. The sync-index was smaller and less variable in Tau- and WT animals. To illustrate this, we estimated a threshold syncindex as the mean+2s.d. of the distribution obtained from the pooled WT animals (which was 1.86). The fraction of epochs exceeding this threshold ranged 1-3% in Tau- animals, and 23-35% in Tau+ animals (Fig 4C). Consequently, the sync-index was on average higher in Tau+ animals than Tau- or WT, at both 5m and 8m (Fig 4D).

Fluctuations in the sync-index may be driven by fluctuations in low frequency power, high gamma power, or both. We therefore analysed the temporal correlation in power at different frequencies (amplitude-amplitude coupling, or AAC (Bruns et al., 2000; Siegel et al., 2012)). For each session (while the animal was at rest) we calculated the temporal correlation between all combinations of frequencies, then averaged across sessions and animals (Fig 5A). Similar frequencies were generally positively coupled – that is, power at similar frequencies modulated in synchrony. Power at very different frequencies was instead usually weakly or even negatively coupled.

**Figure 5 –.**
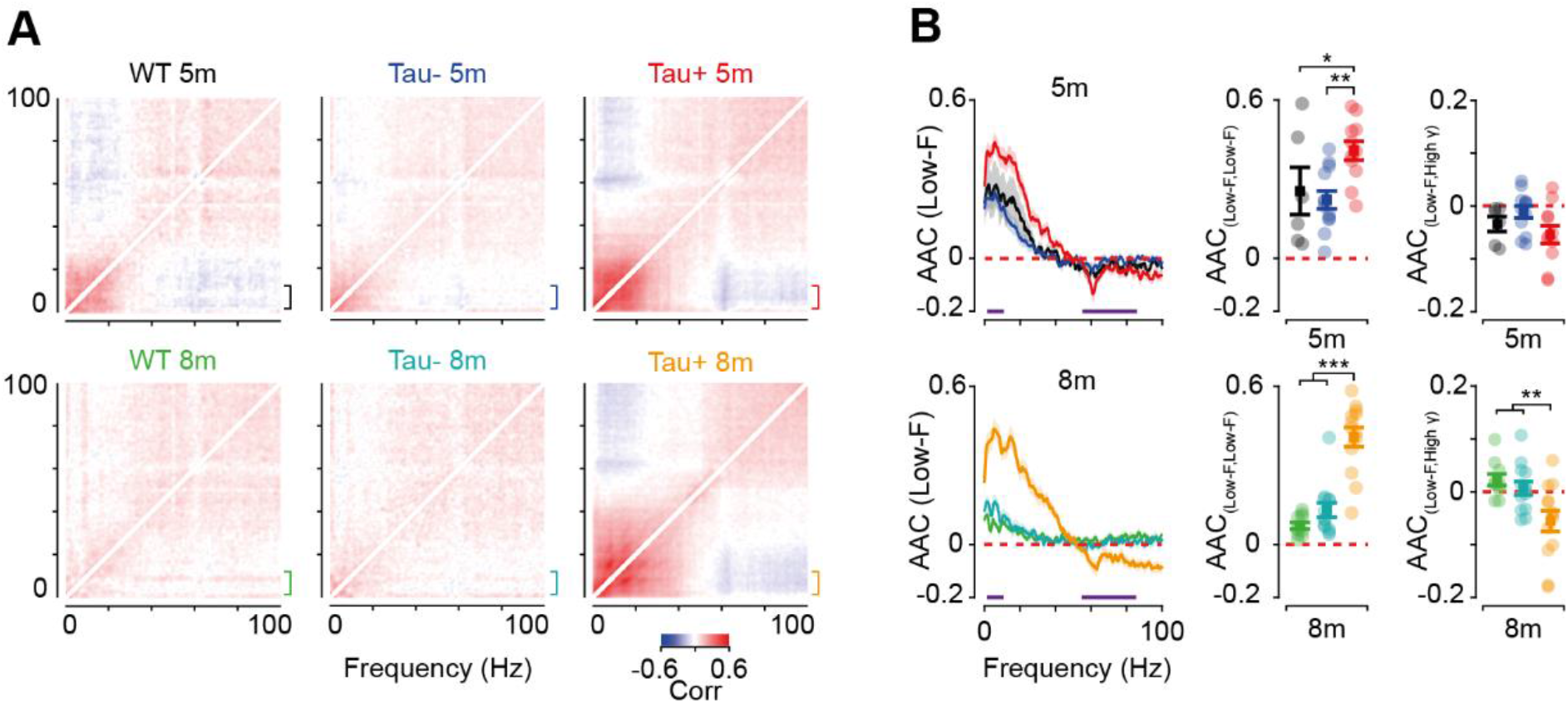
Low frequency activity is more strongly correlated in Tau+ animals during rest periods. (A) Correlation matrices showing the amplitude-amplitude coupling (AAC), a measure of correlation of spectral amplitudes across all frequencies, during rest periods. These revealed stronger amplitude coupling of low frequency rhythms in Tau+ animals at both early and advanced stages of degeneration. Low frequency rhythms also appear to be more negatively correlated with high gamma rhythms in Tau+ animals at both ages. (B) Line plot showing the mean correlation (thick lines) and respective standard error (shaded area) between a low frequency band (2-10Hz) and all other frequencies (indicated by the brackets in A), highlighting the increased correlation of activity within low frequency bands, which are more strongly anti-correlated with high gamma in Tau+ animals (both ranges are indicated by the purple line below the plots). We compared the mean correlation within the low frequencies, and between low and high gamma (55-85Hz) frequency bands across animals and ages during rest periods. Correlation within the low frequency band was larger for Tau+ animals at both ages relative to control animals (WT 5m, 0.255±0.090; WT 8m, 0.072±0.013; Tau- 5m, 0.222±0.034; Tau- 8m, 0.131±0.027; Tau+ 5m, 0.408±0.036; Tau+ 8m, 0.407±0.037; *F*_(2,59)_=28.086; WT x Tau+ 5m, *P*=0.049; Tau- x Tau+ 5m, *P*=0.002; WT x Tau+ 8m, *P*=4.83×10^-8^; Tau- x Tau+ 8m, *P*=8.85×10^-7^). Low frequency and high gamma power were more anticorrelated on average in Tau+ animals (WT 5m, −0.035±0.014; WT 8m, 0.023±0.011; Tau- 5m, −0.012±0.011; Tau- 8m, 0.006±0.013; Tau+ 5m, −0.055±0.018; Tau+ 8m, −0.055±0.020). This difference was significant at more advanced stages of degeneration (*F*_(2,59)_=7.588; WT x Tau+ 8m, *P*=1.64 x10^-3^; Tau- x Tau+ 8m, *P*=8.66×10^-3^).

We found that positive correlation between low frequencies was stronger in Tau+ animals than Tau- or WT animals, at both 5m and 8m (Fig 5B). We also found stronger negative correlation between low frequency and high gamma band power in Tau+ than in either WT or Tau- animals (Fig 5B). The stronger negative correlation suggests that the fluctuations in sync-index reflect anti-phase fluctuations in low- and high frequency power, and not simply fluctuations in one of those bands. The difference between Tau+ and control animals was more pronounced at 8m than at 5m. This primarily reflected a generalised reduction in correlation in WT and Tau- animals, across all frequency bands: Tau+ animals did not show the same age-dependent reduction.

## Discussion

Tau dysfunction is a hallmark of many neurodegenerative diseases, including AD. We measured the effect of tauopathy on visual cortical oscillations in the rTg4510 mouse model, at both early and more advanced stages of tauopathy. Our measurements from visual cortex allowed us to characterise the impact of tauopathy across both low and high frequency rhythms, at different levels of arousal and movement, and in presence and absence of external stimulation. We observed an overall decrease in gamma-power at more advanced stages of tauopathy, and a reduction in stimulus-induced gamma oscillations. Earlier stages of tauopathy, however, were mainly characterized by an increase in low frequency power during resting state. This increase in low frequency power arose because Tau+ animals, but not control animals, intermittently entered states of high synchronisation, in which low frequency oscillations were strongly correlated with each other.

### Impairments of gamma-band activity in tauopathy

We found that gamma power was reduced at more advanced stages of degeneration. This overall reduction in gamma was accompanied by strong reduction in the amplitude of gamma-bumps observed during patterned visual stimulus, but weak if any changes in gamma rhythms observed during presentation of a grey-screen. The different effects of more advanced tauopathy on the two gamma oscillations may be explained by the fact that the two ‘bumps’ appear to have different sources: grey-screen gamma in V1 is thought to be inherited from visual thalamic inputs (Saleem et al., 2017) while stimulus-induced gamma may be generated by local V1 circuits (Veit et al., 2017). Expression of mutant human tau is mainly confined to the neocortex and hippocampus in rTg4510 (Tsien et al., 1996; Santacruz et al., 2005) and is largely absent from visual thalamus. Cortical atrophy and cell death increases rapidly in rTg4510 animals from an age of 6m (Ramsden et al., 2005; Santacruz et al., 2005). By 8m, cortical pyramidal cells show loss of dendritic spines and dendritic atrophy (Crimins et al., 2011), accompanied by loss of axonal boutons (Jackson et al., 2017; Jackson et al., 2020). These morphological changes may result in reduced drive to pyramidal-interneuron networks that are important for the stimulus-induced gamma (Booth et al., 2016b; Chen et al., 2017; Veit et al., 2017). We found little effect of tauopathy on gamma at 5m of age, suggesting that early stages of synaptic dysfunction do not affect these interneuron networks.

### Increased low frequency activity in Tau+ animals during rest

We found that tauopathy had most impact on low frequency oscillations during rest. Low frequency oscillations are a prominent feature of inactive states including anaesthesia, sleep, and quiet wakefulness (Steriade et al., 1993; Greenberg et al., 2008; Poulet and Petersen, 2008; Curto et al., 2009). These synchronized states appear to emerge spontaneously in cortical circuits when there is weak drive to the circuit (Harris and Thiele, 2011; Sanchez-Vives et al., 2017). The increase in low frequency power that we see in tauopathic animals may therefore be explained by a reduced drive to cortical circuits during tauopathy (Spires-Jones and Hyman, 2014; Menkes-Caspi et al., 2015). We find that thalamic inputs to visual cortex (as indexed by narrowband gamma oscillations) appear resilient to tauopathy in these animals. By contrast cortical activity is generally reduced during tauopathy (Busche et al., 2019), and theory suggests that highly synchronised, low frequency states are more prevalent when cortical inhibition is relatively strong (Gao et al., 2017; Chini et al., 2021). The increased low frequency power that we see in tauopathic animals may therefore reflect reduced excitation within visual cortical circuits.

We showed that epochs of synchronization in tauopathic animals are characterized by a large co-fluctuation of activity across a wide range of low frequencies. Fluctuations in delta band (2-4 Hz) were strongly correlated with fluctuations at other low frequencies, up to about 20 Hz, which includes theta and beta bands. Interactions between distal cortical circuits are thought to be primarily carried by oscillations in the theta to beta range (Chen et al., 2017; Fournier et al., 2020; Limanowski et al., 2020). The simpler structure of low frequency oscillations may therefore be consistent with a reduction in coupling (“disconnection” (Delbeuck et al., 2003)) of visual cortex to other cortical circuits. Alternatively, the increased low frequency oscillations may reflect local changes that may in turn also lead to disconnection, by subsuming other rhythms.

We found that epochs of synchronization in tauopathic animals were interleaved with epochs in which cortical activity appeared normal. Most models of cortical function suppose that it operates around a critical point, such that cortex can be driven into and out of synchronised states by small perturbations (Poil et al., 2012). Large increases in low frequency activity may therefore emerge with only a small imbalance in the distribution of excitation and inhibition. The physiological impact of arousal and locomotion is an increase in gamma power and a concomitant decrease in low frequency power (McCormick and Bal, 1997; Vinck et al., 2015). Indeed, we found that movement was sufficient to abolish the differences between Tau+ and control animals. Intermediate states of arousal (large pupil, stationary) also reduced (but did not abolish) some of the differences between tauopathic and control animals. Our observations suggest that locomotion- or arousal-related modulatory inputs to visual cortex may be sufficient to push tauopathic circuits towards, or into, normal activity ranges.

### Relationship to AD and ageing

We found that the impact of tauopathy on low frequency oscillations precedes changes in gamma-range activity. Previous observations in amyloid-β models, also show disruptions to both low frequency and gamma range activity (Scott et al., 2012; Verret et al., 2012; Iaccarino et al., 2016), but whether low frequency disruptions emerge earlier to gamma disruptions is not known. Unlike tau dysfunction, amyloidopathy in known to promote hyperactivity of neuronal ensembles followed by subsequent homeostatic inhibition (Palop et al., 2007), and such imbalances in excitation and inhibition may also lead to increased synchronisation (Verret et al., 2012). Theta rhythms are affected prior to significant amyloid-β dysfunction (Goutagny et al., 2013), and it is therefore possible that impairments to low frequency oscillations also arise before changes in gamma oscillations in amyloid models of AD. We found tauopathic degeneration has more impact on specific components of gamma activity compared to others, and it would therefore be of interest to know if there is a similar disaggregation of gamma-range oscillations in preclinical models of amyloidopathy.

EEG measurements in human patients show that the power of low frequency oscillations is increased in dementia (Coben et al., 1983; Huang et al., 2000), and that these changes are more prominent during resting states (de Haan et al., 2008; Babiloni et al., 2009). The gamma activity that normally accompanies visual stimulation is also impaired in patients of AD (van Deursen et al., 2008). It is striking that the changes that we see in rTg4510 mouse model are so similar to those observed in human patients of dementia and AD. Recent work suggests that modulation of gamma range activity may alleviate progression of AD (Iaccarino et al., 2016; Adaikkan et al., 2019; Chan et al., 2021). In this context, it would be useful to know which components of gamma, if any, better track human AD, as this might offer better therapeutic targets. We also observed that increases in arousal levels, as measured by behavioural variables, reduced the differences in spectral power between tauopathic and control animals. There is some evidence that therapies that reduce apathy and increase motivation improve quality of life (Grasel et al., 2003; Hattori et al., 2011). Better understanding of the relationship between arousal and cortical function in degeneration may therefore offer new opportunities for understanding dementia and AD.

## Acknowledgements

This work was supported by the Medical Research Council grant (R023808) to S.G.S., A.B.S. and Francesca Cacucci., a Sir Henry Dale Fellowship from the Wellcome Trust and Royal Society (200501) to A.B.S, and an International Collaboration Award to S.G.S (with Adam Kohn) from the Stavros Niarchos Foundation / Research to Prevent Blindness.

## Author Contributions

Conceptualization, F.R.R., A.P., A.B.S., and S.G.S.; Methodology, F.R.R., A.P., A.B.S., and S.G.S.; Investigation, F.R.R., A.P., and J.H.; Validation, Formal Analysis, Data Curation, F.R.R., A.P., S.G.S.; Writing – Original Draft, F.R.R., S.G.S, and A.B.S.; Writing – Review & Editing, F.R.R., A.P., J.H., K.G.P., A.B.S., and S.G.S.; Visualization, F.R.R., A.P., A.B.S., and S.G.S.; Funding Acquisition, A.B.S., and S.G.S.; Resources, A.B.S., K.G.P. and S.G.S.; Supervision, A.B.S., and S.G.S.

## Methods

### Animal Experiments

Experiments were performed in accordance with the Animals (Scientific Procedures) Act 1986 (United Kingdom) and Home Office (United Kingdom) approved project and personal licenses. The experiments were approved by the University College London Animal Welfare Ethical Review Board under Project License 70/8637.

In total, 50 double transgenic rTg4510 animals (Ramsden et al., 2005; Santacruz et al., 2005), and 16 wildtype (WT) littermate male mice were obtained at approximately 7 weeks of age from Eli Lilly and Company (Windlesham, UK) via Envigo (Loughborough, UK). In order to suppress mutant tau expression, 25 transgenic mice received doxycycline treatments from 8 weeks of age, including four 10mg/kg bolus oral doses of doxycycline (Sigma) in 5% glucose solution by oral gavage across 4 days, followed by *ad libitum* access to Teklad base diet containing 200ppm doxycycline (Envigo) for the duration of the experiment. These animals were designated as ‘Tau-’, and the remaining 25 transgenic animals which continued to express the tau transgene as ‘Tau+’. WT animals received 4 oral doses of the 5% glucose vehicle solution, followed by *ad libitum* access to standard feed until the end of the experiment. All animals had *ad libitum* access to water. Data from 1 Tau+ animal was removed from the study because histology showed that the recording electrode was placed too deep. Data were obtained in 2 batches and measurements were made in four interleaved cohorts: 2 cohorts of ‘5 months’ (22-26 weeks, 5+7 Tau-, 5+6 Tau+, 2+4 WT); and 2 cohorts of ‘8 months’ of age (31-35 weeks, 6+7 Tau-, 6+7 Tau+, 4+6 WT). All animals were group housed to a maximum of 5 individuals per cage until 3 days before surgery, and then remained singly housed for the remainder of the experiment. Animals were housed under an inverted 12-hour light cycle, and all measurements were carried out during the dark phase of the cycle.

### Surgery

Animals were anaesthetised with 3% isoflurane in a constant flow of oxygen. Preoperative analgesia (Carprieve, 5mg/kg, Norbrook) was administered subcutaneously, and the eyes protected with lubricant ophthalmic ointment (Xailin, Nicox). Anaesthesia was maintained with 1-2% isoflurane in oxygen, and the depth of the anaesthesia was monitored by paw-pinch-withdrawal reflex and breathing rate. Body temperature was maintained using a heating pad. A craniotomy (<1mm^2^) was made over the right primary visual cortex (and a chronic LFP recording electrode (Bear lab chronic microelectrode Monopolar 30070, FHC, USA) was implanted. The coordinates used for all 5m and 8m WT and Tau- mice were 2.8mm lateral, 0.5mm anterior from lambda, and 0.45mm from the cortical surface. Due to degeneration, we adjusted the coordinates for 8m Tau+ mice to 2.7mm lateral, 0.55mm anterior from lambda, and 0.4mm from the cortical surface. A ground screw was implanted over the left prefrontal cortex, and a custom-built stainless-steel metal plate was fixed onto the implant cap. Dental cement (Super-Bond C&B, Sun Medical) was used to cover the skull, ground screw and metal plate, enclosing and stabilizing the electrode. Postoperative analgesic (Metacam, Boehringer Ingelheim, 1mg/kg) was mixed in condensed milk for the animals to consume orally for 3 days after the surgery. Animals were allowed to recover for 7 days before testing started.

### Experimental design

The visual stimulus was a large (80° diameter), sinusoidal grating generated using BonVision (Lopes et al., 2021), presented on a gamma-corrected computer monitor (Iiyama ProLite EE1890SD). The grating (spatial frequency 0.05 cycles/degree), was oriented either −45° or 45° from vertical (counterbalanced within groups), and was presented in a circular aperture with hard edges, outside of which the monitor was held at the mean luminance (‘grey screen’). The grating was modulated in time by a square-wave waveform and reversed contrast (flickered) at a temporal frequency of 0.98 Hz. The display was placed 15cm from – and normal to – the mouse, and centred on the left monocular visual field. The stimulus was warped to maintain visual angle across the monitor.

Mice were habituated to head-fixation over a period of 5 days during which they were secured above a styrofoam wheel and allowed to walk or run while viewing a grey screen. Movement speed was recorded using a rotary encoder; pupil size and position were measured via an infrared camera (DMK 22BUC03, ImagingSource; 30 Hz) focused on the left eye through a zoom lens (Computar MLH-10X Macro Zoom Lens), and extracted from these videos using custom routines in Bonsai (Lopes et al., 2015). Following habituation, animals were tested daily over 9 consecutive days just as described in Papanikolaou et al. (2021). For the purpose of the current study, we only considered recordings made from day 2 to day 8 (7 days total). Each of these recording sessions began with a presentation of a grey screen for 3-5min, followed by the presentation of 10 blocks of the flickering grating, each separated by 30s of grey screen. Each block consisted of 200 reversals, except in 6 of the 5m old animals (2 WT, 2 Tau- and 2 Tau+) where 400 reversals were presented in each block.

LFP signals were acquired, digitised and filtered using an OpenEphys acquisition board (version 1, OpenEphys). The LFP and rotary wheel signals were sampled at 30kHz, and were synchronized with pupil measurements and visual stimulus via the signal of a photodiode (PDA25K2, Thorlabs, Inc., USA) that monitored timing pulses on a small corner of the monitor shielded from the animal. Portions of the same recordings were used to measure visually-evoked potentials reported elsewhere (Papanikolaou et al., 2021). All data have been reanalysed for the current purposes.

### Data analysis

All data were analysed using custom software written in MATLAB (version R2019a, MathWorks, NA).

Wheel speed and pupil diameter signals were mapped to the temporal resolution of spectrograms (described below), and used to define different behavioural epochs for subsequent analysis. Wheel speed was estimated using a quadrature encoder, smoothed with a Gaussian filter (s.d. 50ms). Animals were considered stationary when wheel speed was below 5cm/s. Eye blinks were identified as instances where eye position exceeded the mean plus twice the variance over the recording session, and the corresponding estimates of pupil size were replaced using nearest-neighbour interpolation. Animals were considered to have ‘small pupils’ when pupil diameter was smaller than the mean pupil diameter over the recording session.

LFP signals were filtered with an 8th order Chebyshev Type I low-pass filter and down-sampled to 1kHz using the function *decimate.* These signals were analysed using the Chronux toolbox for MATLAB (Bokil et al., 2010) using the following parameters: tapers, 3,5; window size, 3s; window shift, 1s. Power spectra were obtained by averaging the resultant spectrograms over relevant epochs. Specific oscillations within the high gamma range were characterized with the ‘fitting oscillations & one over f’ (FOOOF) MATLAB toolbox (Donoghue et al., 2020). Normalized spectral power were produced by dividing the power spectrum by FOOOF’s estimate of the aperiodic component at each frequency. The normalized power of gamma range narrowband oscillations was measured by calculating the area-under-the-curve (AUC) of the normalized spectra in a 6Hz window centred on the relevant peak frequency. The parameters used for aperiodic component estimation and peak detection were as follows: peak width limits, [1,8]; max number of peaks, 8; minimum peak height, 0; peak threshold, 3; aperiodic mode, ‘knee’; frequency range, [2, 100].

### Histology

Mice were anaesthetised with isoflurane, and subsequently euthanized with an intraperitoneal overdose of pentobarbital and perfused with 1X phosphate-buffered saline (PBS, Thermo Fisher Scientific). Brains were removed, weighed, and the right hemisphere was fixed in 10% buffered formalin (7-13 months) for immunohistochemistry. Brains were sunk in PBS + 30% sucrose, and 40μm parasagittal sections were cut on a cryostat (Leica CM1520). Sections were heated in a citrate buffer (pH 6.0; Vector Labs) in an oven at 60°C overnight for antigen retrieval. Endogenous peroxidase activity was eliminated by immersing tissue in 0.3% hydrogen peroxide solution (3% in distilled water) for 10 min, followed by washing with PBS with 0.5% Triton X-100. Sections were then incubated with an antibody against phosphorylated-tau (mouse monoclonal AT8, 1:1000, Thermo Fisher Scientific). The Mouse on Mouse Detection Kit (Vector Labs, BMK-2202) was used for subsequent steps as described in the supplied protocol. Tissue was then treated for 5 min with 3,3’-diaminobenzidine (DAB; Vector Laboratories, SK-4105). The slides were coverslipped using Shandon ClearVue Mountant XYL (Thermo Fisher Scientific), and digital images obtained using a Leica Microscope (DMi8 S) coupled with a Leica camera (DFC7000 GT; Leica Microsystems). Greyscale images were processed (blind to genotype/DOX) in Matlab. In each animal, the mean and variance of pixel intensity was estimated across three regions of interest (ROI) of constant area, each on different sections, that encompassed all layers of primary visual cortex.

### Statistical analyses

All data are presented as mean ± standard error of the mean (s.e.m.). All comparisons were performed on SPSS (version 25, IBM). A two-way or a mixed ANOVA design was applied for all comparisons, using phenotype and age as between-subjects factors. In cases where a mixed design ANOVA was used, behavioural variables (e.g. moving) were used as within-subjects factors, and Greenhouse-Geisser corrections for sphericity were applied. All subsequent post-hoc tests were adjusted for multiple comparisons using the Bonferroni correction. Exact *F* statistics with respective degrees of freedom and *P* values for post-hoc comparisons are presented in supplementary tables and figure legends.

